# Contribution of HMGB1 to Keratinocyte inflammation in Recessive Dystrophic Epidermolysis Bullosa

**DOI:** 10.1101/2025.08.11.668806

**Authors:** Kacey Guenther Bui, Ya-Chu Chang, Wannasiri Chiraphapphaiboon, Jianfeng Wang, Christen L. Ebens, Anja-Katrin Bielinsky, Jakub Tolar, Hai Dang Nguyen

## Abstract

Recessive dystrophic epidermolysis bullosa (RDEB) is an inherited skin disorder characterized by fragile skin, blistering, and chronic wounds. Keratinocytes, the primary cells in the epidermis, are directly affected by persistent injury in RDEB, contributing to chronic inflammation. High mobility group box 1 (HMGB1) is elevated in the serum of individuals with RDEB. However, its role in keratinocyte inflammation remains unclear. Here, we report an increase in *HMGB1* expression in keratinocytes at chronic wound sites compared to matched non-wounded skin from an RDEB individual, suggesting a potential link to local inflammation. Pharmacological inhibition of HMGB1 using inflachromene reduced lipopolysaccharide (LPS)-induced inflammatory responses in a keratinocyte cell line, supporting a role for keratinocyte-specific HMGB1 in inflammation. Surprisingly, deletion of *HMGB1* alone or together with its paralogue *HMGB2* did not suppress the inflammatory response to LPS. Furthermore, inflachromene still reduced inflammation in these knockout cells. This unexpected discrepancy between genetic deletion and pharmacologic inhibition points to a more complex role for HMGB1 or off-target effects of the compound. These findings suggest that HMGB1 may contribute to inflammation in keratinocytes, but its exact function needs further investigation.

## INTRODUCTION

Recessive dystrophic epidermolysis bullosa (RDEB) is a rare genetic skin disease caused by mutations in *COL7A1*, the gene coding for type VII collagen. Type VII collagen is an extracellular basement membrane protein that anchors the epidermis to the underlying dermis and plays a key role in maintaining skin integrity. Biallelic mutations in the *COL7A1* gene leads to skin fragility, severe blistering at the dermal-epidermal junction, and chronic wound formation (Cianfarani et al. 2017). While inflammation initially promotes healing, prolonged activation of the inflammatory response by repeated wounding in RDEB further impairs wound resolution. If not resolved, wound chronicity predisposes tissues to the development of metastatic squamous cell carcinoma (SCC), the leading cause of morbidity and mortality in RDEB (Fine et al. 2009). Wound resolution and prevention of carcinogenesis represents an unmet need in the treatment of RDEB, requiring a better understanding of the local inflammatory response within chronic wounds.

Chronic inflammation emerged as a central feature in the pathogenesis of RDEB disease following a seminal study of a monozygotic twin pair with discordant RDEB phenotypes (Odorisio et al. 2014). This study demonstrates the importance of inflammatory signaling via the transforming growth factor beta (TGF-β) axis, providing genetic evidence that links inflammatory response to disease etiology. Evaluation of the microenvironment of chronic skin lesions in RDEB reveals the presence of immune cells, suggesting a pro-inflammatory microenvironment (Riedl et al. 2022). Transcriptomic analysis of skin from three patients with RDEB compared to healthy controls revealed upregulation of genes associated with immune activation, most notably IL-8 and IFN-α-regulated genes (Breitenbach et al. 2015). Pro-inflammatory cytokines are also elevated in the blood of RDEB patients compared to healthy controls (Esposito et al. 2016). In particular, elevated IL-6 serum levels correlates with disease severity (Esposito et al. 2016). While the upregulation of pro-inflammatory signals in RDEB has been clearly demonstrated, cell type-specific contributions to the inflammatory milieu found in the wounds of RDEB patients is not yet fully understood. As the cell of origin for SCC, a better understanding of the specific role keratinocytes play in creating a pro-inflammatory and tumor-permissive environment will enhance our understanding of the pathogenesis of RDEB and development of SCC.

High mobility group box 1 (HMGB1) is ubiquitously expressed across different cell types. In the nucleus, HMGB1 regulates chromatin structure and remodeling, transcription, and DNA replication and repair (Lange and Vasquez 2009). In response to environmental stresses, HMGB1 is released from the cell and acts as a damage-associated molecular pattern (DAMP) to trigger innate immune response and recruit inflammatory cells to damaged sites (Kim et al. 2018; Kim et al. 2020; Kwak et al. 2020). HMGB1 can be actively secreted in response to inflammatory stimuli or passively released from apoptotic cells (Nguyen et al. 2017). In RDEB patients, HMGB1 is elevated locally in wounded skin and systemically in serum relative to healthy controls (Hoste et al. 2015; Petrof et al. 2013; Tamai et al. 2011). Elevated levels of HMGB1 protein have been associated with poor outcomes in multiple inflammatory diseases and malignancies including the RDEB and RDEB-associated SCC (Pandolfi et al. 2016). Keratinocyte-specific HMGB1 primes neutrophils to form neutrophil extracellular trap (NET) in skin wounds, causing a delay in wound healing *in vivo* (Hoste et al. 2019). Instead, NET formation promotes wound-induced tumorigenesis through TNF and RIPK1 kinase activity. Murine studies demonstrate increased keratinocyte HMGB1 expression at wound margins and promotion of keratinocyte and fibroblast activation, angiogenesis, and scar formation by HMGB1 (Dardenne et al. 2013). Despite a link between HMGB1 levels, inflammation, and RDEB disease, the molecular mechanisms of HMGB1 production, secretion, and modulation in keratinocytes remain poorly understood.

In this study, we investigated how HMGB1 contributes to inflammatory responses in keratinocytes. We found that *HMGB1* expression is significantly higher in keratinocytes at wounded regions compared to matched non-wounded region of an RDEB individual. Inhibition of HMGB1 using inflachromene (ICM), an inhibitor that targets HMGB1, reduced LPS-induced inflammation in a keratinocyte cell line. Surprisingly, ICM treatment was still able to suppress LPS-stimulated inflammation in *HMGB1* knockout (HMGB1^KO^) cells. Our findings demonstrate unknown targets of ICM beyond HMGB1 protein and warrant further investigation of HMGB1’s exact function in keratinocyte-specific inflammatory response.

## RESULTS

### RDEB keratinocytes display increased HMGB1 expression

To determine the contribution of *HMGB1* to wound signaling in RDEB, we analyzed single-cell RNA-seq datasets from wounded and matched non-wounded skin biopsies of an RDEB patient who had undergone bone marrow transplantation (Riedl et al. 2022). We focused on four cell clusters: basal keratinocytes, non-basal keratinocytes, myofibroblasts, and fibroblasts (**Figure 1A**), defined according to the expression of cell cluster-specific genes (see Methods). In both basal and non-basal keratinocytes, *HMGB1* expression was significantly higher in wounded compared to non-wounded skin (**Figures 1B-D**). In contrast, HMGB1 expression in fibroblasts was modestly reduced in wounded relative to non-wounded skin and remained unchanged in myofibroblasts regardless of wounding (**Figures 1E-F**). These results suggest a potential link between keratinocyte-specific response to chronic wounding in RDEB and *HGMB1* expression.

**Figure 1.**
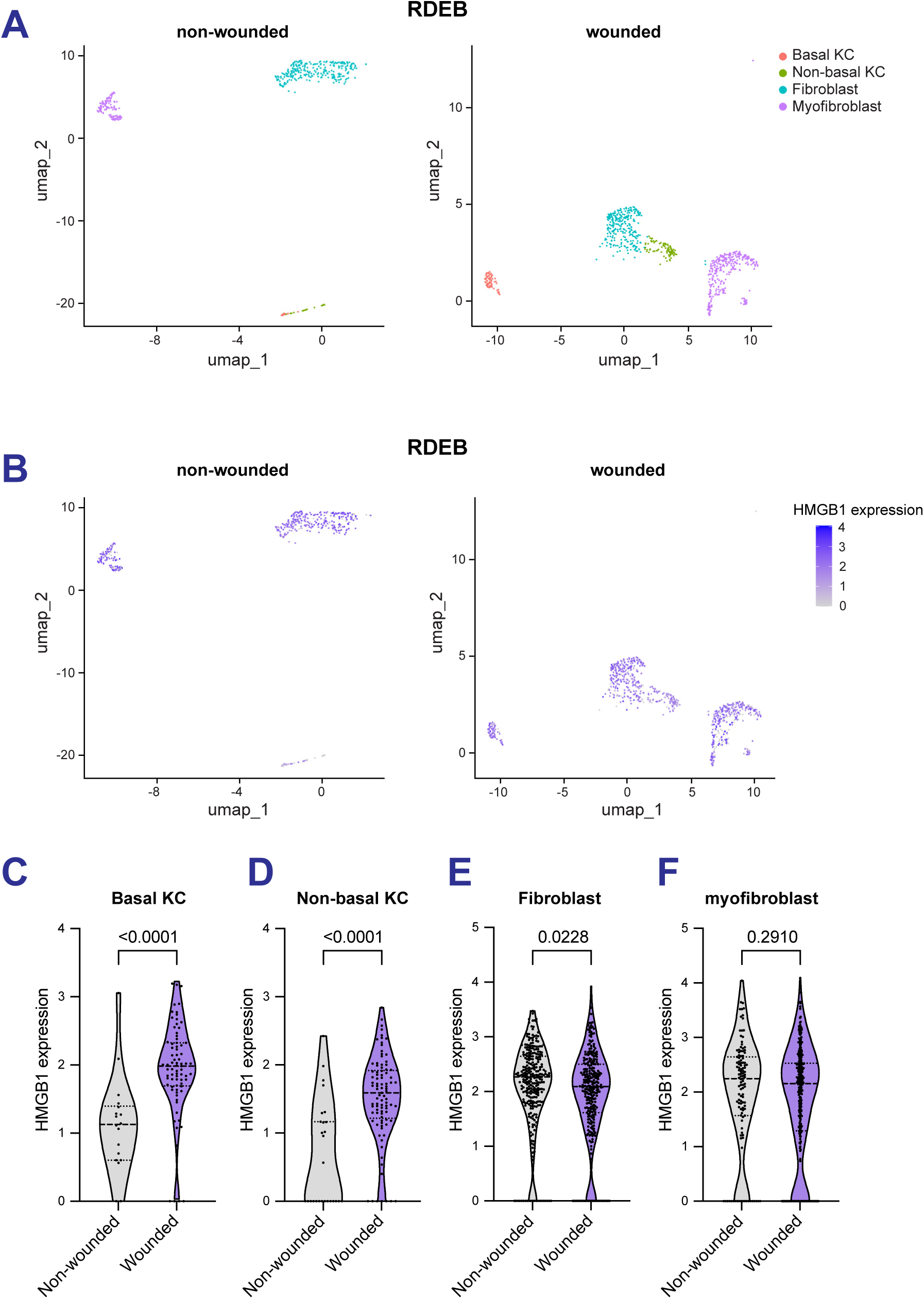
Increased *HMGB1* expression in keratinocyte cells at a wounded site of an RDEB patient. **(A-B)** Single-cell RNA sequencing analysis of keratinocyte and fibroblast cells from wounded and non-wounded skin obtained from an RDEB patient. Identification of epidermal cell clusters by gene expression (see methods) is shown in A. Expression of *HMGB1* in each cluster is shown in B. **(C-F)** Violin plots of *HMGB1* expression in indicated cell types from wounded and non-wounded skin from an RDEB patient. Each dot represents *HMGB1* expression per nucleus. Statistical analysis was performed using unpaired Student’s t-test.

### Pharmacologic inhibition of HMGB1 suppresses LPS-induced inflammatory response in keratinocytes

The increase in *HMGB1* expression in keratinocytes prompted us to investigate how HMGB1 regulates inflammation. To address this, we treated the human keratinocyte cell line N/TERT-2G cells with LPS for 24 hours in the presence or absence of HMGB1 inhibitor ICM. Cell supernatants were collected and assessed for the release of cytokines and chemokines. Among the 71 cytokines and chemokines tested, IL-6, TNFα, GROα (CXCL1), IL-8 (CXCL8), and platelet-derived growth factor (PDGF)-AA/AB isoforms were significantly elevated upon LPS stimulation and were suppressed by ICM treatment (**Figures 2A-C**, **Supplemental Table S1**). Although IL-1α, MIG/CXCL9, and IL-18 were also significantly induced by LPS, they were not affected by HMGB1 inhibition (**Figures 2A**, **Supplemental Table S1**). We independently validated LPS-induced IL-6 release in both N/TERT-2G and a second human keratinocyte cell line, HEK001 (**Figures 2D-E**) as a marker of inflammation in keratinocytes. Notably, ICM treatment also suppressed LPS-induced IL-6 in both cell lines. Taken together, these results suggest that HMGB1 affects a subset of inflammatory factors in response to LPS stimulation in keratinocytes.

**Figure 2.**
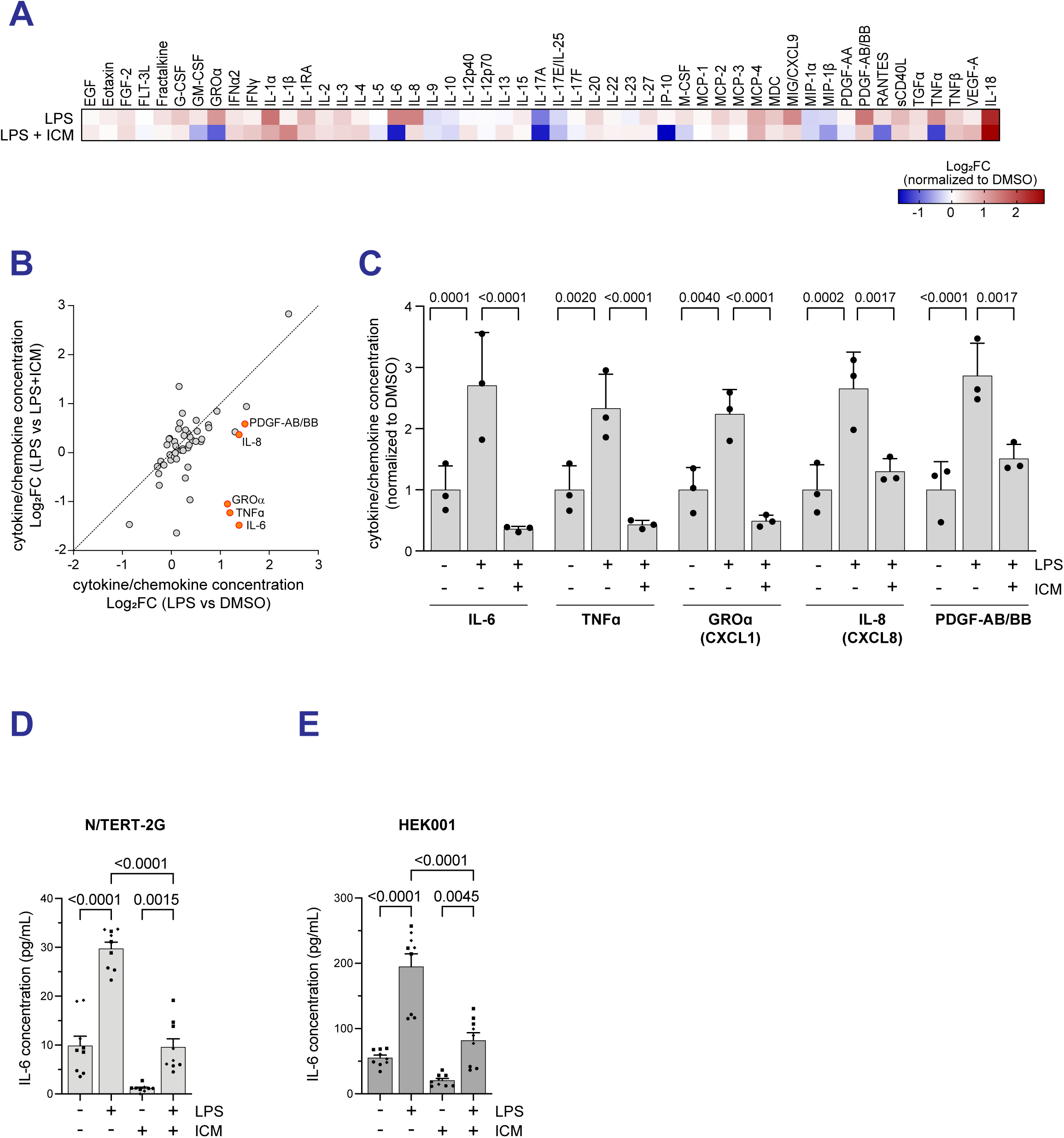
Pharmacologic inhibition of HMGB1 suppresses LPS-induced inflammation response keratinocyte cell line. **(A-C)** Average fold-change of indicated cytokines/chemokines following LPS treatment in the presence or absence of inflachromene, ICM from 3 biological replicates. N/TERT-2G cells were pre-treated with 5 µM ICM or vehicle for 3 hours prior to vehicle or 100 µg/mL LPS treatment for an additional 24 hours. Samples were normalized to vehicle-treated samples with the average of 3 biological replicates depicted in the heatmap in A. A two-dimensional plot of average fold-change of individual cytokine/chemokines from 3 biological replicates is shown in B. Orange-filled dots illustrate factors that were significantly induced by LPS and suppressed in LPS+ICM-treated condition. A bar graph demonstrating fold-change of cytokine/chemokine release of indicated factors normalized to vehicle-treated sample is depicted in C. Individual points represent independent biological replicates. One-way ANOVA statistical analysis was separately performed for each factor. **(D-E)** ELISA analysis of supernatant IL-6 concentration in N/TERT-2G (D) and HEK001 (E) cells using IL-6 ELISA kit (n=3 biological replicates denoted by shape, each with 3 technical replicates). Cells were treated with 10 μM (N/TERT-2G) or 5 μM (HEK001) ICM or vehicle for 24 hours prior to vehicle or 20 μg/mL LPS treatment for an additional 24 hours. One-way ANOVA statistical analyses were performed.

### HMGB1 pharmacologic inhibition, but not *HMGB1* deletion, suppresses LPS-induced IL-6 stimulation in keratinocytes

To further assess the regulation of HMGB1-dependent inflammation in keratinocytes, we assessed the impact of a *HMGB1* gene deletion using the CRISPR/Cas9 strategy. Successful *HMGB1* deletion, both in pooled populations and single-cell clones, was confirmed by TIDE analysis (Brinkman et al. 2014) (**Supplemental Figure S1A**). HMGB1 protein levels were confirmed by immunoblotting and immunofluorescence using an HMGB1 specific antibody (**Figure 3A**, **Supplemental Figure S1B**). Karyotype analyses of HMGB1^KO^ single clones and a HGMB1^WT^ clone that had undergone the same CRISPR/Cas9 editing revealed similar karyotypes between HMGB1^WT^ and HMGB1^KO^ cells (**Supplemental Figure S1C**). Both wildtype and knockout HMGB1 cells had gained one copy each of chromosomes 7 and 20 at the end of the clone isolation process.

**Figure 3.**
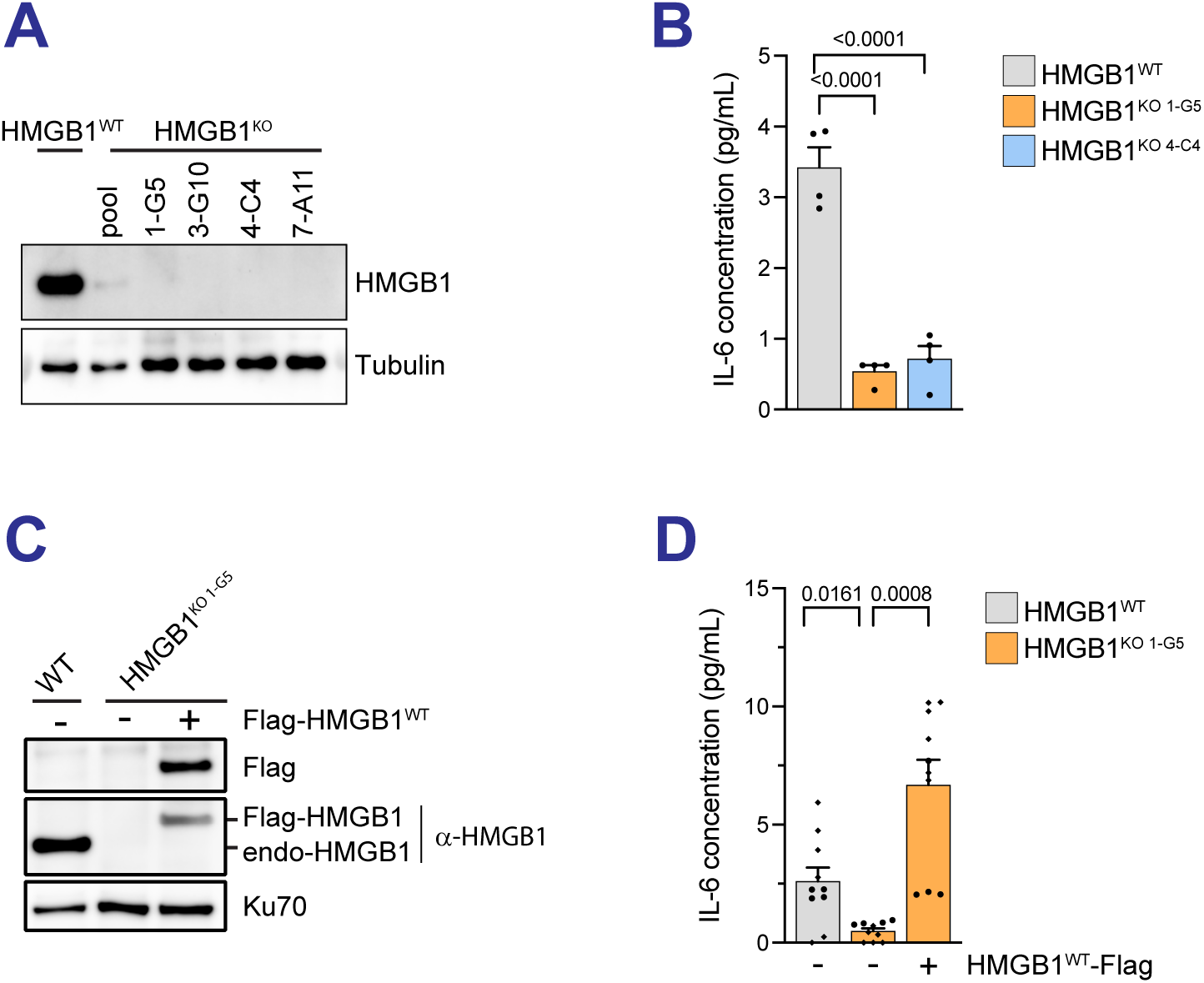
HMGB1 regulates IL-6 release at basal conditions. **(A)** Validation of HMGB1^KO^ N/TERT-2G cells by western blot. **(B)** ELISA analysis of supernatant IL-6 concentration of HMGB1^WT^ and two independent HMGB1^KO^ cells. **(C)** Immunoblot analysis of endogenous and exogenous Flag-HMGB1 expression in indicated cell lines. **(D)** ELISA analysis of supernatant IL-6 concentration at basal levels in indicated cells.

To validate the role of HMGB1 in inflammation, we first assessed IL-6 release under basal conditions in HMGB1^WT^ and two independent HMGB1^KO^ clones. HMGB1^KO^ cells showed reduced IL-6 release compared to isogenic HMGB1^WT^ cells, suggesting that HMGB1 contributes to inflammation under basal conditions (**Figure 3B**). Overexpression of a three-Flag, N-terminal-tagged HMGB1 (Flag-HMGB1) in HMGB1^KO^ cell restored IL-6 release (**Figures 3C-D**). This was confirmed in a second HMGB1^KO^ clone (**Supplemental Figure S1D**). These results confirm that IL-6 production is dependent on HMGB1 in the absence of other inflammatory stimuli like LPS.

Next, we evaluated the role of HMGB1 under LPS-stimulated conditions. In contrast to the basal state, HMGB1^KO^ cells exhibited elevated IL-6 release following LPS treatment, comparable to that of HMGB1^WT^ cells (**Figure 4A**, compare lanes 2 vs 5). Moreover, treatment with ICM still effectively suppressed LPS-induced IL-6 release in both HMGB1^WT^ and HMGB1^KO^ cells (**Figure 4A**, compare lanes 2 vs 3 and lanes 5 vs 6). These results indicate that the LPS-induced IL-6 release is independent of HMGB1 yet remains sensitive to ICM treatment.

**Figure 4.**
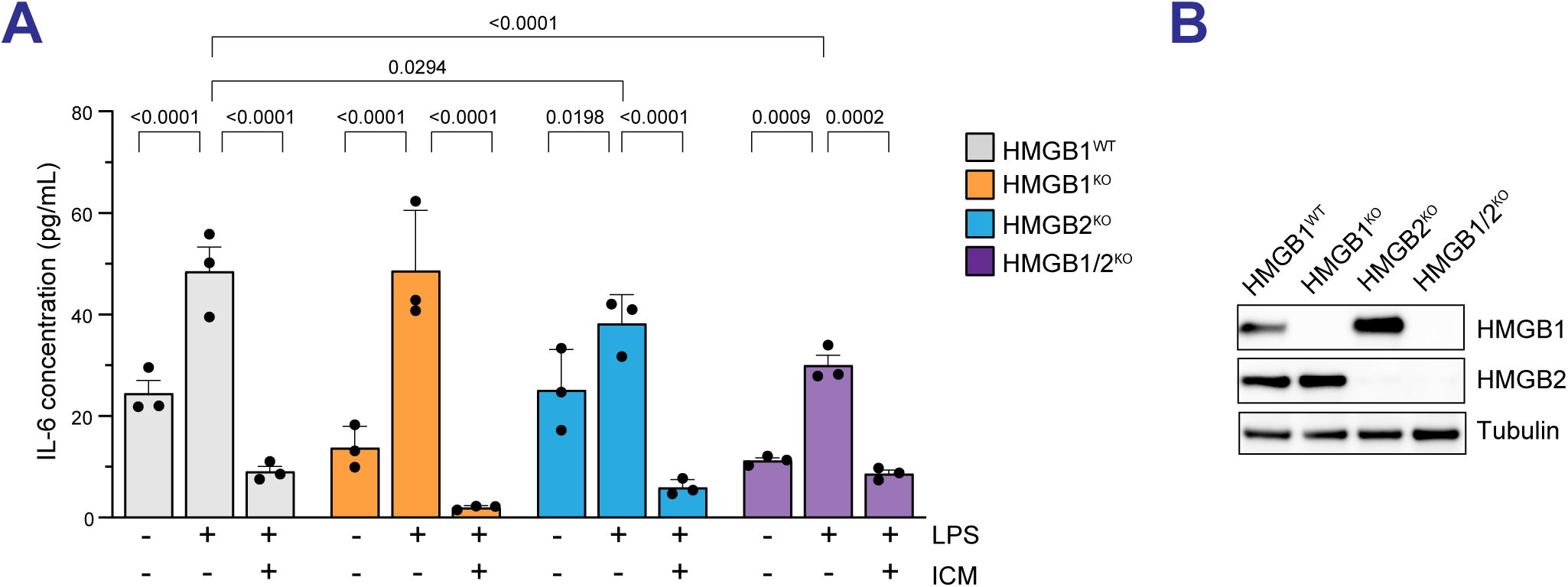
Pharmacologic inhibition of HMGB1, but not *HMGB1* knockout, suppresses LPS-induced IL-6 release. **(A)** Asynchronous cells were pre-treated with 5 µM ICM or vehicle for 3 hours, followed by 100 µg/mL LPS or vehicle treatment for an additional 24 hours. Supernatants were collected for IL-6 release analysis by ELISA. **(B)** Immunoblot analysis of HMGB1 and HMGB2 levels in indicated cell lines.

The ICM compound inhibits both HMGB1 and HMGB2 (Lee et al. 2014). We hypothesized that HMGB2 may compensate for the loss of HMGB1. To test this, we deleted the *HMGB2* gene in both HMGB1^WT^ and HMGB1^KO^ cells and confirmed successful knockout by western blot (**Figure 4B**). In contrast to HMGB1^KO^, HMGB2^KO^ showed modest reduction in IL-6 secretion compared to LPS-treated HMGB1^WT^ or HMGB1^KO^ cells (**Figure 4A**, lanes 2 and 5 vs lane 7). ICM still significantly suppressed LPS-stimulated IL-6 secretion in HMGB2^KO^ cells (**Figure 4A**, lane 9). Similar to HMGB1^KO^, HMGB1/2 double knockout cells (HMGB1/2^KO^) also maintained IL-6 responsiveness upon LPS treatment, and ICM continued to suppress this response (**Figure 4A**). While the absolute IL-6 levels in LPS-stimulated HMGB1/2^KO^ cells were significantly lower than in LPS-stimulated HMGB1^WT^ cells, the fold-change induction of LPS-to-DMSO treatment was not significantly different (HMGB1^WT^: 1.993 vs HMGB1/2^KO^: 2.70, p-value > 0.99). We excluded comparisons with HMGB1^KO^ cells for fold-change induction analysis due to their reduced basal IL-6 levels. Taken together, our results demonstrate that pharmacologic inhibition of HMGB1 using ICM, but not genetic deletion of *HMGB1* or *HMGB2*, suppressed LPS-induced IL-6 release. These findings raise the intriguing possibility that ICM may target additional factors beyond HMGB1/2 proteins to regulate inflammation in keratinocytes.

## DISCUSSION

In this study, we investigated the role of HMGB1 in regulating IL-6 secretion in keratinocytes, aiming to clarify its function in the context of RDEB. Single-cell RNA sequencing of wounded and non-wounded skin regions from an RDEB patient revealed increased *HMGB1* expression specifically in keratinocytes, suggesting a role in RDEB-associated skin injury and inflammation. Through profiling a panel of cytokines and chemokines, we found that pharmacologic inhibition using ICM suppressed a subset of LPS-stimulated inflammatory markers, including IL-6. This suggests that HMGB1 selectively mediates certain inflammatory responses in keratinocytes. However, HMGB1^KO^ keratinocyte cells retained a functional inflammatory response to LPS stimulation that could be suppressed by ICM. The discrepancy between pharmacologic inhibition and genetic deletion warrants further investigation to precisely define the role of HMGB1 in keratinocyte-specific inflammation.

Although HMGB1 levels are elevated in both the serum and wounded skin regions of RDEB patients with chronic blistering (Hoste et al. 2015; Petrof et al. 2013; Tamai et al. 2011), its molecular function in keratinocytes remains unclear. A previous study showed that keratinocyte-specific deletion of *HMGB1* delayed wound healing and promoted SCC tumorigenesis *in vivo* (Hoste et al. 2019), highlighting a potential role for HMGB1 in the progression from chronic wounds to carcinogenesis. Importantly, this did not apply to myeloid cells. Our observed increased *HMGB1* expression in keratinocytes at wounded sites in an RDEB patient also suggests a keratinocyte-specific, intrinsic inflammatory response that may contribute to disease pathogenesis. In addition to this keratinocyte-specific response, single-cell RNA sequencing of the same patient also identified distinct immune cell populations within the wounded microenvironment (Riedl et al. 2022), indicating that both intrinsic (keratinocyte-driven) and extrinsic (immune cell-derived) inflammatory signals may contribute to RDEB pathogenesis. Future studies investigating the interplay between these inflammatory components will be critical for developing new therapeutic strategies.

HMGB1 is constitutively expressed across many cell types. While its role in immune cell– mediated inflammation is well established, its function in keratinocytes is less well understood. Our finding that *HMGB1* expression is upregulated in keratinocytes from wounded compared to non-wounded skin suggests a potential role for both context- and cell type-specific HMGB1 function in RDEB disease etiology and warrants further investigation. Due to the higher expression of *HMGB1* in fibroblast cells, the differential *HMGB1* expression may not be as drastic compared to keratinocyte cells. Although ICM has been used as an HMGB1 inhibitor, we observed that it fully suppresses inflammation even in HMGB1^KO^ cells, implying that ICM may target additional inflammatory mediators beyond HMGB1 and HMGB2. A recent study identified that ICM also targets Kelch-like ECH-associated protein 1 (Keap1), a negative regulator of Nuclear factor erythroid 2-related factor 2 (Nrf2) in an antioxidant response pathway (Yim et al. 2024). It will be interesting to determine how Keap1 contributes to a keratinocyte-specific inflammatory response. Identifying ICM-specific targets in keratinocytes may open new therapeutic strategies to dampen inflammation as a potential treatment for RDEB patients. Finally, the HMGB1^KO^ and HMGB2^KO^ keratinocyte lines developed in this study provide a valuable platform for dissecting keratinocyte-specific HMGB1/2-mediated inflammatory signaling and validating future HMGB1/2-targeted pharmacologic inhibitors.

## MATERIALS AND METHODS

### Cell Lines

N/TERT-2G keratinocyte cells were obtained from Dr. Rheinwald through the laboratory of Ellen van den Bogaard at Radboud University Medical Centre in Nijmegen, the Netherlands (Dickson et al. 2000; Smits et al. 2017). HEK001 human epidermal keratinocyte cells were obtained from ATCC (CRL-2404) (Sugerman and Bigby 2000). Both cell lines were grown in Keratinocyte Serum-Free Medium (K-SFM; Gibco 17005-042) with 50 units/mL penicillin and 50 μg/mL streptomycin (Gibco 15070-063). N/TERT-2G cell media was supplemented with epithelial growth factor (EGF; Gibco 10450-013, 0.2 ng/mL), bovine pituitary extract (BPE; Gibco 13028-014, 25 μg/mL), and 0.4 mM final concentration of CaCl2 (Teknova C0477). HEK001 cell media was supplemented with EGF (Gibco 10450-013, 5 ng/mL). N/TERT-2G cells were passaged using TrypLE Express Enzyme (Gibco 12604-210), and HEK001 cells were passaged using 0.25% Trypsin-EDTA (Gibco 25200-056). Cells were pelleted at 300-350 x g for 5 min, resuspended into single-cell suspension, and counted using a Vi-Cell BLU cell viability analyzer/automated cell counter. Cells were re-plated at ∼10,000-17,000 cells/cm^2^ to maintain optimal cell growth. Media was changed every 2 days and cells were passaged every 3-4 days (<70% confluent) to maintain cells in logarithmic growth phase. Mycoplasma detection was performed every 6-12 months using a commercially available detection kit (EZ-PCR^TM^ Mycoplasma Detection Kit, Sartorius 20-700- 20 or Venor™ GeM Mycoplasma Detection Kit, MilliporeSigma MP0025).

For Flag-HMGB1-expressing cells, supernatant containing virions was collected at 72 hours post transfection and added to N/TERT-2G HMGB1^KO^ cells in the presence of polybrene (10 μg/mL, Millipore TR-1003-G) by the spinoculation method. Cells expressing Flag-HMGB1 were selected in media containing 16 μg/mL blasticidin (A1113903) for 4 days. Flag-HMGB1 expression was confirmed by immunoblotting. Flag-HMGB1-expressing cells were maintained in media containing 4 μg/mL blasticidin.

### Generation of knockout cell lines by CRISPR/Cas9

Synthetic sgRNAs targeting *HMGB1* (GAUACUCACGGAGGCCUCUU) and *HMGB2* (AAAAAUUACGUUCCUCCCAA) genes were designed using Synthego CRISPR design tool. sgRNAs and Cas9 mRNA were purchased from Synthego. Cas9 mRNA and sgRNA were nucleofected into low passage N/TERT-2G cells by electroporation (Neon^TM^ transfection system) using optimized settings for N/TERT-2G cells (1500V, 10ms, 3 pulses) and for HEK001 cells (1400V, 20ms, 2 pulses). To increase cell survival during single cell cloning, 96-well tissue culture plates were coated with VitroCol Type I collagen (Advanced Biomatrix 5007) according to the manufacturer’s protocol. Cells were plated at 1-5 cells per well in “conditioned media” (media from cells cultured 24-48 hours at optimal density in logarithmic growth phase, sterile filtered and mixed 1:1 with fresh media). Cell stocks of edited clones were frozen in freezing media (50% K-SFM, 20% DMEM/Ham’s F12, 20% FBS, 10% DMSO).

To determine the efficiency of CRISPR-induced insertions and deletions (indels), genomic regions flanking CRISPR/Cas9 cut sites were amplified by PCR and sequenced by Sanger sequencing. Primers used to screen *HMGB1-KO* at exon 3 are Forward: 5’- ATTCAGAGCAGACTCGGGCGGA-3’ and Reverse: 5’-TGTGATGCATTGGACAGGGTGC- 3’. The resulting amplicons are analyzed using a decomposition algorithm called TIDE that identifies indels present in the cell population given a known cut site. P-values generated during TIDE analysis were used to determine the significance of the detected indels (Brinkman et al. 2014).

### HMGB1 cDNA design and lentiviral production

The wild-type *HMGB1* coding sequence with 3xFLAG-N-terminal tag (HMGB1-Flag) flanked by attB1/2 sequences was synthesized and cloned into pUC57 cloning vector by Gene Universal. The cDNA was serially cloned into pDONOR221 and subsequently swapped into pLenti6.2/V5-DEST Lentiviral expression vector by Gateway cloning. Successful cloning of the 3Flag-HMGB1 cDNA insert was confirmed by Sanger sequencing.

For lentiviral production, HEK293T cells were co-transfected with 10µg of each plasmid: pLenti6.2-Flag-HMGB1, pCMV-dR8.2 dVPR (Addgene #8455), and pCMV-VSV-G (Addgene #8454) using the calcium phosphate-mediated ProFection Mammalian Transfection System (Promega, E1200) for 72 hours.

### Karyotyping

N/TERT-2G cell lines were verified approximately every year by karyotyping. Following 3.0 colcemid treatment, 20 metaphase spreads were analyzed, and karyotypes were generated according to standard cytogenetic protocol.

### Cell plating and treatments

Logarithmically growing cells were used for all experiments. 75-100 x 10^3^ cells/well were plated in 24-well plate in K-SFM media, respectively. After allowing cell attachment for overnight, cells were replenished with new K-SFM media and pre-treated with 5 μM (HEK001) or 10 µM (N/TERT-2G) inflachromene (Sigma 533060) or 5 μM ICM (Cayman Chemicals 17006) or vehicle control followed by 100 µg/mL Lipopolysaccharides (LPS) from *E. Coli* strain O55:B5 (Sigma L6529) or O111:B4 (Sigma L2630; Lot 0000369272) for 24 hours prior to collection. Different durations of ICM treatment will be noted for specific experiments.

### Enzyme-Linked Immunosorbent Assay (ELISA)

Supernatants were collected, centrifuged at >10,000x*g* and used immediately (or stored at -80°C for ≤ 3 months). The ELISA kit was purchased commercially (ELISA MAX^TM^ Deluxe Set Human IL-6, BioLegend 430516). The protocol was followed according to the manufacturer except for IL-6 protein standards, which were resuspended in cell culture media and supernatants were incubated overnight at 4°C to optimize signal. Supernatant samples were run in duplicate and read using an M1000 (Tecan). Analysis was performed in GraphPad prism using a 2^nd^ order polynomial (quadratic) nonlinear regression to interpolate a standard curve.

The 71-Plex array of human cytokine/chemokine as shown in Figure 2 was done by Eve Technologies (Cat #HD71). Supernatants were done in biological triplicates. Raw cytokines/chemokines calculation are reported in **Supplemental Table S1**. For Figure 2A, the fold-change difference in LPS-treatment alone or combined with ICM was normalized to respective vehicle treated samples.

### Immunoblots

Whole-cell lysates were prepared in 1% sodium dodecyl sulfate (SDS) lysis buffer or radioimmunoprecipitation (RIPA) assay buffer separated on Bolt^TM^ 4-12% Bis-Tris gradient gels (Bio-Rad NW04122), transferred onto 0.45 μM polyvinylidene fluoride (PVDF) membranes. Clarity Western enhanced chemiluminescence (ECL) (Bio-Rad 1705060) or WesternBright Quantum (VWR 103254-878) chemiluminescent substrate was used to detect bound alkaline phosphatase. Signal was detected using a ChemiDoc Imaging System (Bio-Rad). Antibodies used: HMGB1 [EPR3507] (Abcam #ab79823), HMGB2 (D1P9V, CST #14163), FLAG^®^ M2 (Sigma #F1804), α-Tubulin (DM1A, Sigma #T9026), Ku70 (Genetex #GTX70271), anti-mouse (VWR, 102646-160) and anti-rabbit IgG F(c) γ Goat Polyclonal Antibody conjugated HRP (VWR, 102645-182).

### Immunofluorescence

Cells were cultured on cover slides and fixed using 3% sucrose/2% PFA for 15 min at room temperature (RT) and subsequently permeabilized using 0.25% Triton-X100 for 5 min on ice. Slides were incubated in blocking buffer (10% milk/3% BSA in PBS containing 0.1% Triton-X100) for 1 hour at RT prior to staining overnight at 4°C with HMGB1 antibody (EPR3507, Abcam #ab79823) diluted 1:500 in blocking buffer. Slides were washed three times with PBS containing 0.1% Tritan-X100 pre- and post-secondary antibody staining for 1 hour at RT using Alexa Fluor® 488 Donkey Anti-Rabbit IgG (H+L) (Jackson ImmunoResearch 711-545-152) diluted 1:1000 in blocking buffer. Nuclei were counterstaining using DAPI (Sigma D9542) and mounted using ProLong^TM^ Gold Antifade Mountant (Invitrogen P36930) prior to imaging using the 60X objective on a Leica DMi8 fluorescence microscope.

### Statistical Analysis

Data was analyzed via t-test (for comparison of two conditions), one-way ANOVA (for single variable data sets) or two-way ANOVA (for multiple variable data sets) followed by Tukey’s Honest Significant Difference (HSD) post-hoc test to adjust for multiple comparisons; p < 0.05 was considered statistically significant.

### Single-cell RNA sequencing analysis

Single-cell RNA sequencing dataset was previously reported (Riedl et al. 2022). The workflow for obtaining scRNA-seq gene expression results was implemented using R version 4.4.2 and the Seurat package version 5.2.1 (Hao et al. 2024). Initial quality control filtering was applied to each single-cell dataset, retaining only cells with more than 200 and fewer than 2500 detected RNA features (genes), and mitochondrial gene expression percentages below 20%. After filtering, each dataset underwent a standardized preprocessing pipeline, including data normalization, identification of highly variable features, scaling, and dimensionality reduction through principal component analysis (PCA) with 19 principal components. Subsequently, clustering was performed using a resolution of 0.7 (Riedl et al. 2022), and cells were visualized via UMAP projections based on these principal components. Cell identities were determined manually based on established marker gene sets: Keratinocytes were subdivided into basal and non-basal populations, distinguished by differential expression of keratin and epidermal differentiation markers (Basal KC: K15+, KRT1+, KRTDAP+, KRT10+, S100A7+; Non-basal KC: K15-, KRT1+, KRTDAP+, KRT10+, S100A7+). Fibroblasts (FB) were identified by the expression of extracellular matrix and mesenchymal markers (“COL1A1”, “COL1A2”, “COL3A1”, “COL6A1”, “COL6A2”, “VIM”, “DCN”, “LUM”, “FBLN1”, “PDGFRA”) (Solé-Boldo et al. 2020), while myofibroblasts (MyoFB) were defined through expression of smooth muscle-related and matrix genes (“ACTA2”, “TAGLN”, “MYH11”, “MYL9”, “CNN1”, “COL1A1”, “COL3A1”, “PDGFRB”) (Buono et al. 2023; Guerrero-Juarez et al. 2019; Gur et al. 2022; Valenzi et al. 2019).

## Ethics Statement

No conflict of interest was reported by the authors.

## Data Availability

All relevant data generated or analyzed during this study are included in the published article. Further information and requests for resources and reagents should be directed to Drs. Hai Dang Nguyen, Jakub Tolar, Anja-Katrin Bielinsky.

## Acknowledgements

We thank Dr. Rheinwald for N/TERT-2G cells and Molly Lynch at the Cytogenetic studies in the University of Minnesota Cancer Genomics Shared Resource (CGSR) of the Masonic Cancer Center at the University of Minnesota. K.G.B is supported by the National Institutes of Health (NIH) National Cancer Institute Predoctoral Individual National Research Service Grant Award (F31 CA281039). K.G.B. was partially supported by the National Institutes of Health’s National Center for Advancing Translational Sciences, grants TL1R002493 and UL1TR002494. The content is solely the responsibility of the authors and does not necessarily represent the official views of the National Institutes of Health’s National Center for Advancing Translational Sciences. Y.C. was supported by the Targets of Cancer Training Program (NIH T32CA009138). H.D.N. is supported by grants from the Masonic Cancer Center, Edward P. Evans Foundation Discovery Research Grant, the National Heart, Lung, and Blood Institute (R01HL163011), and the 2022 AACR Career Development Award to Further Diversity, Equity, and Inclusion in Cancer Research, which is supported by Merck, grant number 22–20–68-NGUY. AKB was supported by NIH R35GM141805. J.T was supported by NIH/NIAMS R01-AR063070. The authors would like to thank the University of Minnesota Foundation for their continued support. This research was funded in part through the NIH/NCI Cancer Center Support Grant P30 CA008748.

## Author Contributions

Conceptualization – KGB, JT, AKB, HDN; Data curation: KGB, WC, YCC, JW, CE, HDN; Formal analysis: KGB, WC, YCC, JW, HDN; Investigation: KGB, WC, YCC, JW, CE, HDN, JT, AKB; Supervision: JT, AKB, HDN; Writing, review and editing: all authors.

---

**Supplemental Figure S1.**
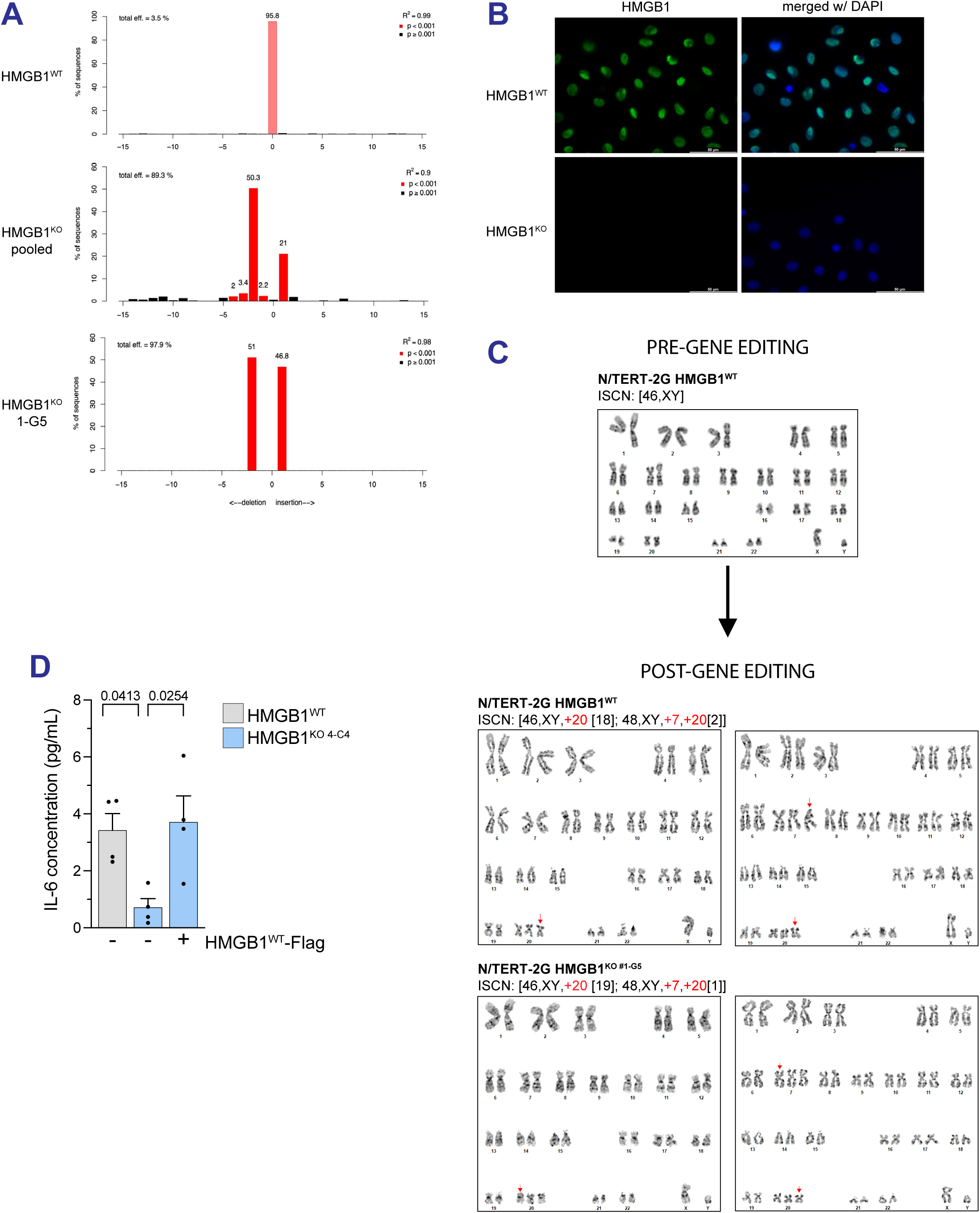
Generation and validation of HMGB1^KO^ N/TERT-2G cells. **(A-B)** Validation of HMGB1^KO^ N/TERT-2G cells by TIDE analysis in (A) and immunofluorescence (in B). **(C)** Representative karyotype images of HMGB1^WT^ and HMGB1^KO^ N/TERT-2G cells. **(D)** ELISA analysis of supernatant IL-6 concentration at basal levels in indicated cells.

**Supplemental Table S1.** Cytokine/Chemokine analysis in keratinocytes. Cytokine and chemokine concentration from a 71-plex array panel were analyzed from 3 biological replicates. Data points were used to generate Figures 2A-B.

